# A high-quality genome assembly of the North American Song Sparrow, *Melospiza melodia*

**DOI:** 10.1101/850990

**Authors:** Swarnali Louha, David A. Ray, Kevin Winker, Travis Glenn

**Affiliations:** Institute of Bioinformatics, University of Georgia, Athens, GA, USA; Department of Biological Science, Texas Tech University, Lubbock, TX, USA; University of Alaska Museum, Fairbanks, AK, USA; Department of Environmental Health Science, College of Public Health, University of Georgia, Athens, GA, USA

**Keywords:** *Melospiza melodia*, Dovetail genomics, Passeriforms, whole genome sequencing, *de novo* assembly

## Abstract

The song sparrow, *Melospiza melodia*, is one of the most widely distributed species of songbirds found in North America. It has been used in a wide range of behavioral and ecological studies. This species’ pronounced morphological and behavioral diversity across populations makes it a favorable candidate in several areas of biomedical research. We have generated a high-quality *de novo* genome assembly of *M. melodia* using Illumina short read sequences from genomic and *in vitro* proximity-ligation libraries. The assembled genome is 978.3 Mb, with a coverage of 24.9×, N50 scaffold size of 5.6 Mb and N50 contig size of 31.7 Kb. Genes within our genome assembly are largely complete, with 87.5% full-length genes present out of a set of 4,915 universal single-copy orthologs present in most avian genomes. We annotated our genome assembly and constructed 15,086 gene models, a majority of which have high homology to related birds, *Taeniopygia guttata* and *Junco hyemalis*. In total, 83% of the annotated genes are assigned with putative functions. Furthermore, only ~7% of the genome is found to be repetitive; these regions and other non-coding functional regions are also identified. The high-quality *M. melodia* genome assembly and annotations we report will serve as a valuable resource for facilitating studies on genome structure and evolution that can contribute to biomedical research and serve as a reference in population genomic and comparative genomic studies of closely related species.

## INTRODUCTION

The oscine passerines (Order Passeriformes) are songbirds having specialized vocal learning capabilities. Many species of songbirds have been widely used by neuroscientists to study the processes underlying memory and learning and social interactions (Doupe et al. 1999, White 2010). The song sparrow (*Melospiza melodia*) is one of the most morphologically diverse songbirds found in North America, with 26 recognized subspecies (Pruett et al. 2008). It has been recognized as a model vertebrate species for field studies of birds and has been the subject of extensive research integrating behavioral and ecological studies over the last 70 years (Arcese et al. 2002). Though several species of songbirds have been sequenced and studied (Warren et al. 2010, Jarvis et al. 2014), few offer the plethora of biomedical research potential presented by the song sparrow. For example, differences in neural growth and song-center brain development among different populations of song sparrows could play an instrumental role in basic research involving human brain neurogenesis (NIH 2001). The molecular processes underlying the regeneration of “hair” cells in the auditory system of song sparrows might also provide therapies useful in human hearing loss (Hawkins et al. 2003, Hawkins and Lovett 2004). Several other areas of biomedical research in which the song sparrow might serve as a model system are hepatic lipogenesis (Gosler 1996, Schubert et al. 2007), craniofacial development (Brugmann et al. 2010, Powder et al. 2012), and variations in body size (Sutter et al. 2007, Allen et al. 2010). The last of these is a polygenic trait, and elucidation of the underlying gene network affecting different metabolic pathways can help clarify several biological phenomena, including human diseases. Given its significant biomedical potential and experimental tractability in the field and aviary, the song sparrow will continue to be used for answering research questions related to mechanisms causing variation in behavior, morphology, and demographics across populations (Arcese et al. 2002, Nietlisbach et al. 2015).

Prior work on population genetics of song sparrows in Alaska has shown how sequential bottlenecking of populations in the Aleutian Archipelago has given rise to a naturally inbred population with large body size (Pruett and Winker 2005). Previous work has also been done on the song sparrow transcriptome, developing genomic markers to screen at population levels (Srivastava et al. 2012). A high-quality genome assembly of *M. melodia* furthers the development of genomic markers to screen loci associated with phenotypic traits of interest. An ever-growing number of songbirds have sequenced genomes, but relatively few have been published so far, including the American crow (*Corvus brachyrhynchos*), golden-collared manakin (*Manacus vitellinus*; Jarvis et al. 2014), Zebra finch (*Taeniopygia guttata*; Warren et al. 2010), medium ground finch (*Geospiza fortis*; Parker et al. 2012) and the dark-eyed junco (*Junco hyemalis*; Friis et al. 2018). In this study, we provide the genome assembly of *Melospiza melodia*, a member of the family Passerellidae; this will facilitate genomic and phylogenetic comparisons with other songbirds, and will also serve as a reference genome for comparative genomic studies.

Our high-quality draft genome assembly of *M. melodia* was created by combining both traditional Illumina paired-end libraries and a *de novo* proximity-ligation Chicago library. The Chicago library method together with Dovetail Genomics’ HiRise software pipeline is designed to significantly reduce gaps in alignment arising from repetitive elements in the genome (Putnam et al. 2016) and increases assembly contiguity. The draft genome was annotated using transcribed RNA and protein sequences from *M. melodia* and related songbird species, *Junco hyemalis* and *Taeniopygia guttata*. Genomic features of interest other than coding sequences, such as microsatellites, repeat elements, transposable elements, and non-coding RNA, were also annotated and the genome assembly was evaluated for quality by comparing it to related avian species.

## METHODS

### Library preparation and *de novo* shotgun assembly

The *de novo* assembly of the song sparrow genome was constructed using Illumina paired end libraries. Genomic DNA was isolated from a blood sample of a single male song sparrow, obtained from the wild in the Aleutian Islands of Alaska on 16 Sep 2003 and archived as a voucher specimen at the University of Alaska Museum (http://arctos.database.museum/guid/UAM:Bird:31500). We sheared the genomic DNA using the Covaris S2 (Covaris, Woburn, MA, USA) targeting a 600bp average fragment size. The sheared DNA was end-repaired, adenylated, and ligated to TruSeq LT adapters using a TruSeq DNA PCR-Free Library Preparation Kit (Illumina, San Diego, CA, USA). We purified the ligation reaction using a Qiaquick Gel Extraction Kit (Qiagen, Venlo, The Netherlands) from a 2% agarose gel. We sequenced the library on an Illumina HiSeq 2500 at the HudsonAlpha Institute for Biotechnology (Huntsville, AL, USA) to obtain paired-end (PE) ~100 bp reads. The sequence data consisted of 276 million read pairs sequenced from a total of 41.3 Gbp of paired-end libraries (~50× coverage). Reads were trimmed for quality, sequencing adapters, and mate pair adapters using Trimmomatic (Bolger et al. 2015). The reads were assembled at Dovetail Genomics (Santa Cruz, CA, USA) using Meraculous 2.0.4 (Chapman et al. 2011) with a *k-mer* size of 29. This yielded a 972.4 Mbp assembly with a contig N50 of 22.5 Kbp and a scaffold N50 of 33 Kbp.

### Chicago library preparation and scaffolding the draft genome

To improve the *de novo* assembly, a Chicago library was prepared at Dovetail Genomics using previously described methods (Putnam et al. 2016). Around 500 ng of high-molecular-weight genomic DNA (mean fragment length = 50 kbp) was used for chromatin reconstitution *in vitro* and fixed with formaldehyde. Fixed chromatin was digested with *Dpn*II, the 5’ overhangs filled in with biotinylated nucleotides, and free blunt ends were ligated together. After ligation, crosslinks were reversed and DNA was purified from protein. Purified DNA was treated to remove biotin that was not internal to ligated fragments. Next, DNA was sheared to ~350 bp mean fragment size and sequencing libraries were generated using NEBNext Ultra enzymes (New England Biolabs, Ipswich, MA, USA) and Illumina-compatible adapters. Biotin-containing fragments were isolated using streptavidin beads before PCR enrichment of the library. The Chicago library was sequenced on an Illumina HiSeq 2500 to produce 47 million 150 bp paired end reads (1-50 kb pairs).

Dovetail Genomics’ HiRise scaffolding software pipeline (Putnam et al. 2016) was used to map the shotgun and Chicago library sequences to the draft *de novo* assembly using a modified SNAP read mapper (http://snap.cs.berkeley.edu). The separations of Chicago read pairs mapped within draft scaffolds were analyzed by HiRise to produce a likelihood model for genomic distance between read pairs, and the model was used to identify and break putative misjoins, to score prospective joins, and make joins above a threshold. After scaffolding, shotgun sequences were used to close gaps between contigs.

### Identification of microsatellites and transposable elements

Transposable elements (TEs) in the song sparrow genome were identified using a combination of *de novo* and homology-based TE identification methods, in addition to a manual curation step (Platt et al. 2016). First, we used RepeatModeler v1.0.11 (Smit and Hubley 2008-2015) with default parameters (File S1) to generate a custom repeat library consisting of 672 consensus repeat sequences. RepeatModeler uses two *de novo* repeat identification programs, RECON v1.08 (Bao and Eddy 2002) and RepeatScout v1.0.6 (Pevzner et al. 2005), for identifying repetitive elements from sequence data. To ensure accurate and complete representation of putative TEs, the RepeatModeler derived consensus sequences were filtered for size (>100 bp), and then subjected to iterative homology-based searches against the genome, followed by manual curation (Platt et al. 2016). The final set of manually curated TEs was queried against CENSOR (Kohany et al. 2006) and TEclass (Abrusan et al. 2009) for classification. TEs not identifiable in CENSOR were also searched against the NCBI nucleotide and protein databases using BLASTN and BLASTX respectively. Finally, a custom *de novo* repeat library comprising song sparrow-specific TEs and existing simple and low complexity repeats in birds was used to screen for repeats in the song sparrow genome assembly with RepeatMasker v4.0.9.

Microsatellites in the song sparrow genome were identified and described with GMATA v2.01 (Wang et al. 2016) with sequence motifs ranging in length from 2-20 bp, and each motif repeated at least 5 times (File S2).

### *De novo* gene annotation and function prediction

Genes were predicted in the song sparrow genome with the MAKER v2.31.9 genome annotation pipeline (Campbell et al. 2014). A custom repeat library of 900 repeat sequences consisting of TEs identified in the song sparrow genome and other existing avian repeat elements was used to soft mask the genome. Transcriptome evidence sets for MAKER included the assembled song sparrow transcriptome (Srivastava et al. 2012) and Trinity (v2.4.0) mRNA-seq assemblies from multiple tissues of *Junco hyemalis* (Peterson et al. 2012, NCBI BioProject Accession: PRJNA256328). Protein evidence sets used by MAKER included annotated proteins for song sparrow, *Junco hyemalis*, and *Taeniopygia guttata* from the NCBI Protein database. The MAKER pipeline consisted of the following steps: 1) Transcriptomic and protein evidence sets were used to make initial evidence-based annotations with MAKER; 2) the initial annotations were used to train two *ab initio* gene predicters: Augustus (Stanke et al. 2006), which was trained once, and SNAP (Korf 2004), which was iteratively trained twice; and 3) the trained gene prediction tools SNAP and Augustus were used to generate the final set of gene annotations (File S3-S8).

Functional annotations of the predicted genes were obtained by making homology-based searches with BLASTP against the Uniprot/Swiss-Prot protein database (Pundir et al. 2016, File S9). InterProScan v5.29 (Zdobnov et al. 2001) was used to find protein domains associated with the genes. The putative functions and protein domains were added to the gene annotations using scripts provided with MAKER (File S9).

To quantitatively assess the completeness of the song sparrow genome assembly and annotated gene set, we ran BUSCO (Benchmarking Universal Single-Copy Orthologs) v3.0.2 (Waterhouse et al. 2017) with 4,915 single-copy orthologous genes in the Aves lineage group (Aves_odb9; https://busco.ezlab.org/), using “chicken” as the Augustus reference species (File S10). The 4,915 orthologous genes are present in at least 90% of the 40 species included within the Aves lineage group, and thus are likely to be found in the genome of related species. We also used the JupiterPlot pipeline (https://github.com/JustinChu/JupiterPlot) to visually compare the zebra finch (*T. guttata*) genome assembly (Warren et al. 2010) to our assembly in a Circos plot, using the largest scaffolds making up 95% of our genome assembly, and all scaffolds greater than 100 kbp in the Zebra finch genome (File S11). The zebra finch was selected because it was the first fully sequenced songbird genome, it is one of the most complete bird genomes, and it is often used for comparative genomic studies in birds.

### Non-coding RNA prediction

Transfer RNAs (tRNAs) were predicted in the song sparrow genome with tRNAscan-SE v2.0 (Lowe and Chan 2016, File S12). A training set comprising eukaryotic tRNAs was used to train the covariance models employed by tRNAscan-SE, and tRNAs were searched against the genome with Infernal v1.1.2 (Nawrocki et al. 2014). tRNAscan-SE also provides functional classification of tRNAs based on a comparative analysis using a suite of isotype-specific tRNA covariance models. A random sample of 10 predicted tRNAs were selected and searched against the tRNA databases GtRNAdb (Chan and Lowe 2016) and tRNAdb (Juhling et al.2009).

Identification of miRNAs (microRNAs), snoRNAs (small nucleolar RNAs), snRNAs (small nuclear RNAs), rRNAs (ribosomal RNAs), and lncRNAs (long non-coding RNAs) was achieved by using a homology-based prediction method. Structural homologs to eukaryotic ncRNA covariance models from the Rfam database v14.1 (Gardner et al. 2009) were searched against the song sparrow genome using Infernal’s “cmscan” program (File S13). All low-scoring overlapping hits and hits with an E-value greater than 10^-5^ were discarded, and the remaining ncRNAs were grouped into different classes.

Lastly, we compared the predicted classes of different ncRNAs in the song sparrow genome to those reported in the genomes of related birds, *Taeniopygia guttata* and *Ficedula albicollis* (collared flycatcher).

## DATA AVAILABILITY

Raw reads have been deposited in the NCBI Sequence Read Archive (SRR10491484 and SRR10451714 for the Meraculous assembly, and SRR10424475 for the Chicago HiRise assembly). Supplemental File S1 contains submission script for RepeatModeler. Supplemental File S2 contains primary configuration file used to run GMATA (default_cfg.txt). Supplemental File S3 contains submission script for MAKER. Supplemental File S4 contains MAKER executable file (maker_exe.ctl). Supplemental File S5 contains specifications for downstream filtering of BLAST and Exonerate alignments (maker_bopts.ctl). Supplemental File S6 contains primary configuration of MAKER specific options (maker_opts.ctl). Supplemental File S7 contains scripts for training SNAP. Supplemental File S8 contains scripts for training Augustus. Supplemental File S9 contains scripts for running BLASTP and InterProScan for functional annotation of predicted genes; and scripts for adding the functional annotations to gene annotation files. Supplemental File S10 contains submission script for BUSCO. Supplemental File S11 contains submission scripts for JupiterPlot pipeline. Supplemental File S12 contains submission script for tRNAscan-SE. Supplemental File S13 contains submission script for Infernal. Supplemental File S14 contains classification of predicted transposable elements. Supplemental File S15 contains annotation of microsatellites with their genomic locations. Supplemental File S16 contains percentage of different microsatellites present in the genome. Supplemental File S17 contains frequency of occurrence of microsatellites in each scaffold of the genome. Supplemental File S18 contains the distribution of the length of microsatellites. Supplemental File S19 contains predicted function of annotated genes by BLASTP. Supplemental File S20 contains prediction of protein domains, GO annotations and pathway annotations of predicted genes by InterProScan. Supplemental File S21 contains sequence and structure of tRNAs identified in the Song Sparrow genome. Supplemental File S22 contains classification of predicted tRNAs. Supplemental File S23 contains classification of different ncRNAs predicted in the genome with Infernal. Supplemental File S24 contains the M. melodia Chicago HiRise genome sequence (Mmel_1.0). Supplemental File S25 contains annotations for the M. melodia genome assembly. Supplemental Table S1 contains genome sizes of birds related to M. melodia. Supplemental Figure S1 contains the distribution of the percentage of annotated genes with their corresponding AED scores.

## RESULTS AND DISCUSSION

### Assembly

We produced the *de novo* genome assembly of song sparrow, with a total length of 978.3 Mb, using a Chicago library and the HiRise assembly pipeline. The N50 scaffold size was 5.6 Mb and contig size was 31.7 Kb. This assembly showed significant improvement over the initial shotgun assembly, with a 169-fold increase in scaffold N50 and a 60-fold increase in scaffold N90 (Table 1). These increases in scaffold size were also accompanied by an increase in assembly contiguity, with the total number of scaffolds decreasing from 74,832 to 13,785 (Fig 1, Table 1).

**Table 1:** A comparison of assembly quality statistics from the initial shotgun sequencing assembled by Meraculous and the final HiRise assembly.

|  | **Meraculous Assembly** | **Chicago HiRise Assembly** |
| --- | --- | --- |
| Total length | 972.4 Mb | 978.3 Mb |
| Scaffold N50 | 33 kb | 5.58 Mb |
| Scaffold N90 | 5 kb | 303 kb |
| Scaffold L50 | 7,552 scaffolds | 48 scaffolds |
| Scaffold L90 | 35,731 scaffolds | 324 scaffolds |
| Longest scaffold | 366,149 | 26,942,064 |
| Number of scaffolds | 74,832 | 13,785 |
| Number of scaffolds > 1kb | 74,806 | 13,768 |
| Contig N50 | 22.5 kb | 31.7 kb |
| Number of gaps | 53,577 | 95,490 |
| Percent of genome in gaps | 1.427% | 1.847% |

**Figure 1:**
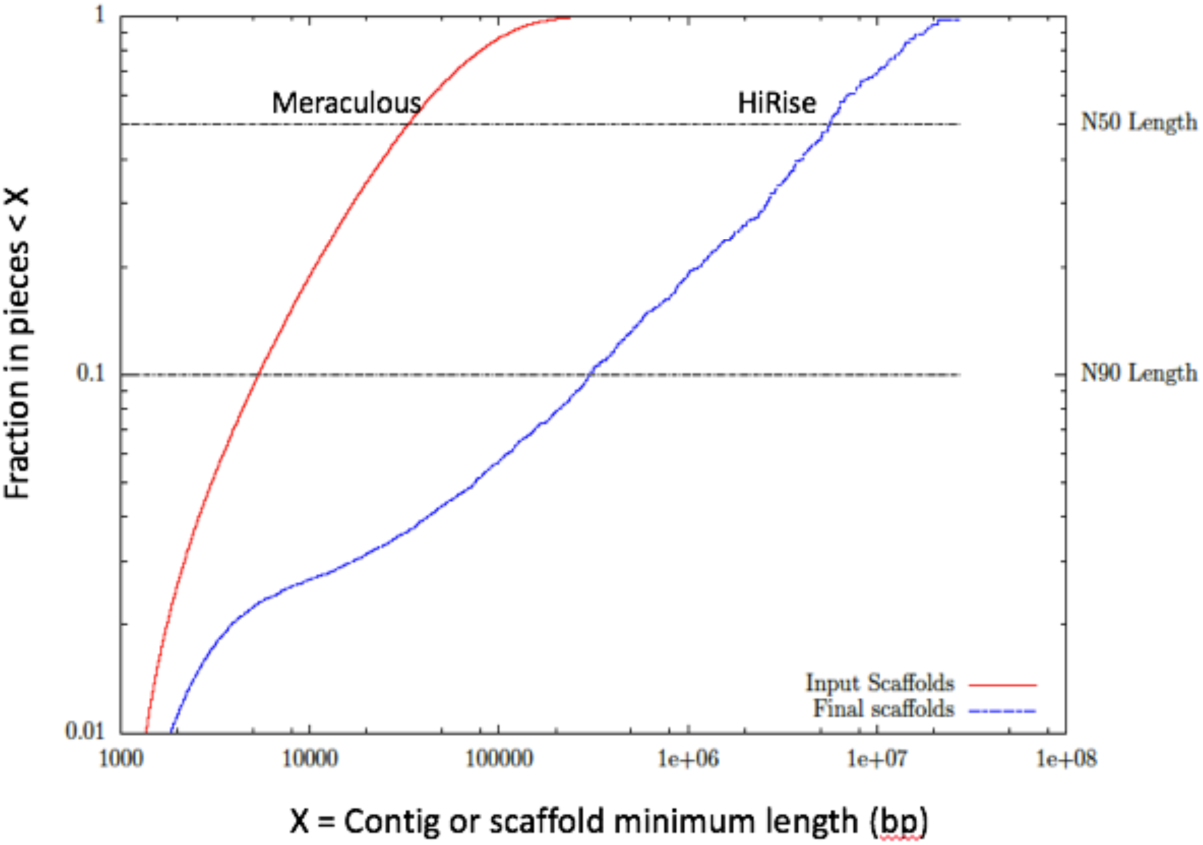
Comparison of assembly contiguity.

### Microsatellites and Transposable Elements

In total, 89 as yet unnamed TEs were identified in the song sparrow genome. Fifty-five of these did not have any significant matches in CENSOR (Kohany et al. 2006) and are considered novel (File S14). A TE was considered to have a significant match to a known element in CENSOR only when it had a length of at least 80 bp and 80% identity to the known element over 80% of its length, the 80-80-80 rule (Schulman et al. 2007). The predicted TEs were classified into DNA transposons and retrotransposons (i.e. LINEs, LTRs, and SINEs) using CENSOR and TEclass (File S14). Approximately 7.4% of the genome comprises repeats with the majority of that consisting of TEs (~ 48%). Among the different TEs, LTRs (~ 40%) and LINEs (~ 49%) were found to be most abundant (Table 2). The song sparrow genome assembly was found to be less repetitive when compared to sequenced genomes of related songbirds, primarily due to the lower content of LTRs and LINEs than other songbirds (Fig 2).

**Table 2:** Number and percentage of repeats in the *M. melodia* genome assembly.

| **Classification** | **Number of copies** | **Percentage of assembly** |
| --- | --- | --- |
| LINEs | 104,032 | 3.01 |
| LTRs | 85,276 | 2.83 |
| SINEs | 6,695 | 0.08 |
| DNA Transposons | 13,521 | 0.21 |
| Unclassified | 4,884 | 0.12 |
| **Total transposable elements** | **214,408** | **6.25** |
| Satellites | 569 | 0.00 |
| Low complexity repeats | 38,561 | 0.20 |
| Microsatellites | 192,996 | 0.90 |
| **Total** | **446,534** | **7.35** |

**Figure 2:**
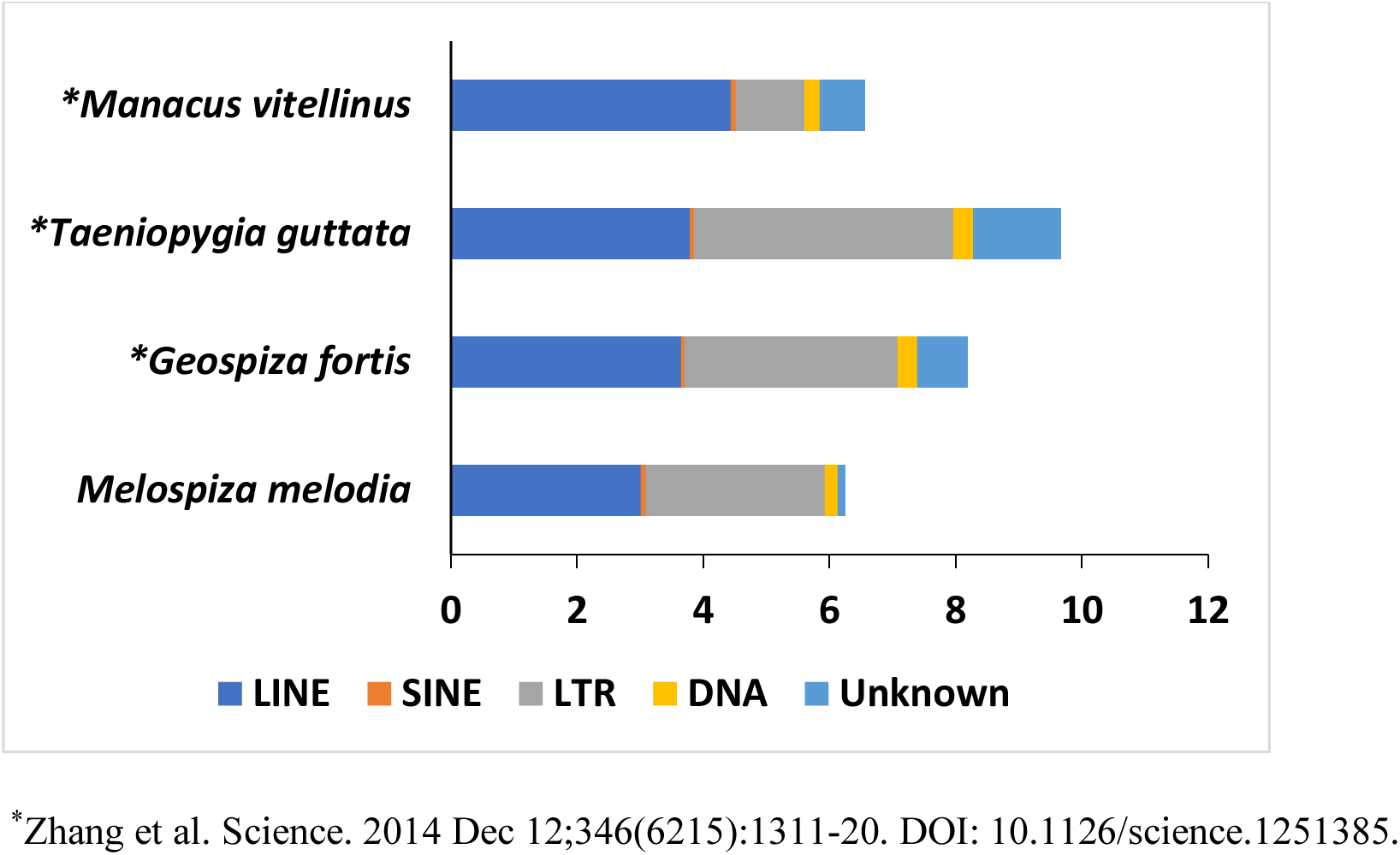
Comparison of percentages of transposable elements (TEs) among related songbirds.

Overall, 112,419 microsatellites with motifs ranging in size from 2-20 bp were found in the song sparrow genome (File S15 contains all microsatellites with their genomic locations). The majority of the microsatellites were made up of 2-, 3-, 4-, and 5-mers, with 2-mers making up about 71% of all microsatellites identified (Fig 3, File S16). The distribution of the top base-pair composition of microsatellite motifs present in the genome is shown in Fig 4. The frequency of occurrence of microsatellites in every scaffold and a distribution of their lengths are provided in Files S17 and S18, respectively.

**Figure 3:**
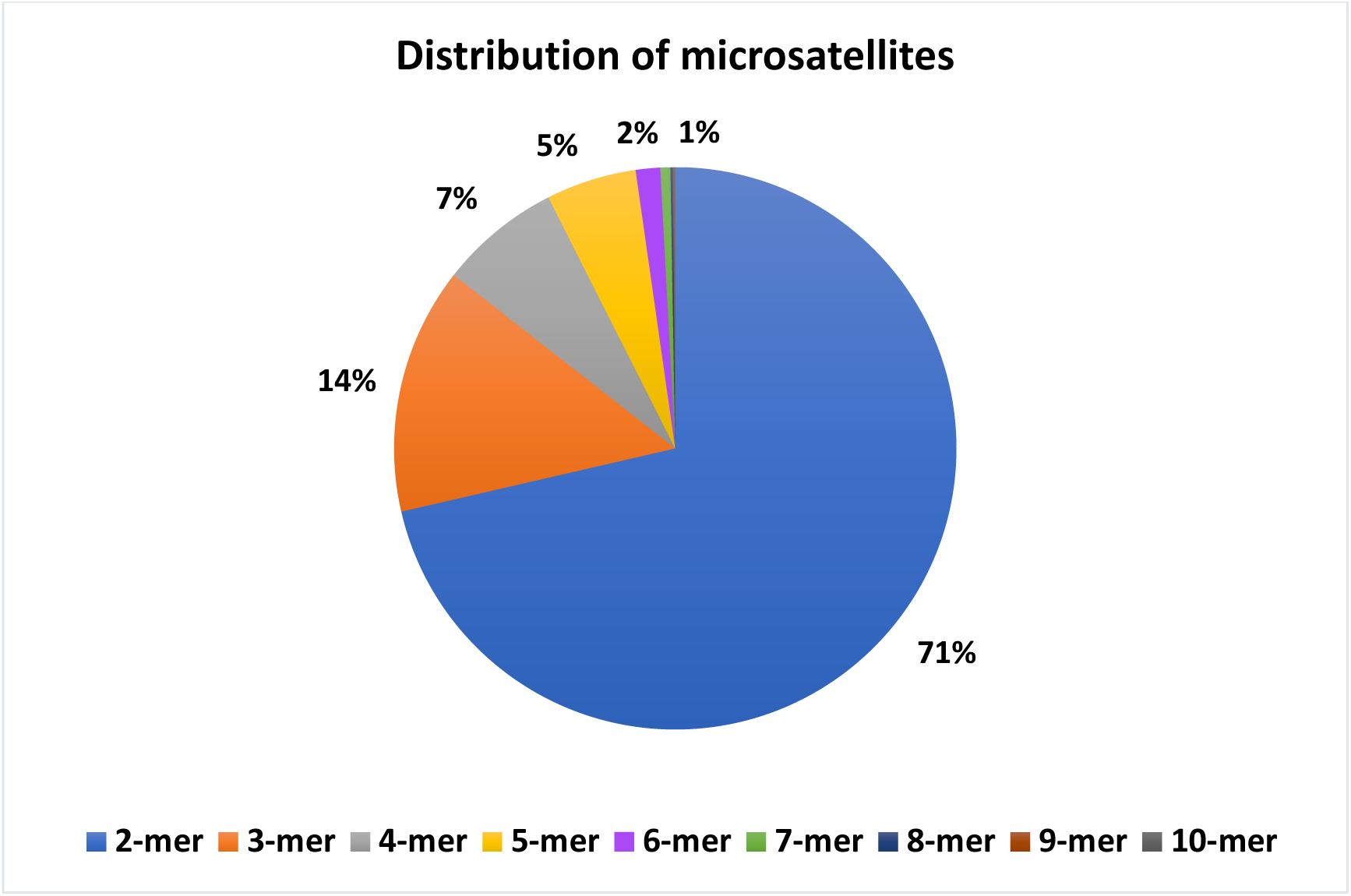
Distribution of microsatellite repeat motif size classes in the *M. melodia* genome assembly (details are given in Supplemental File S15).

**Figure 4:**
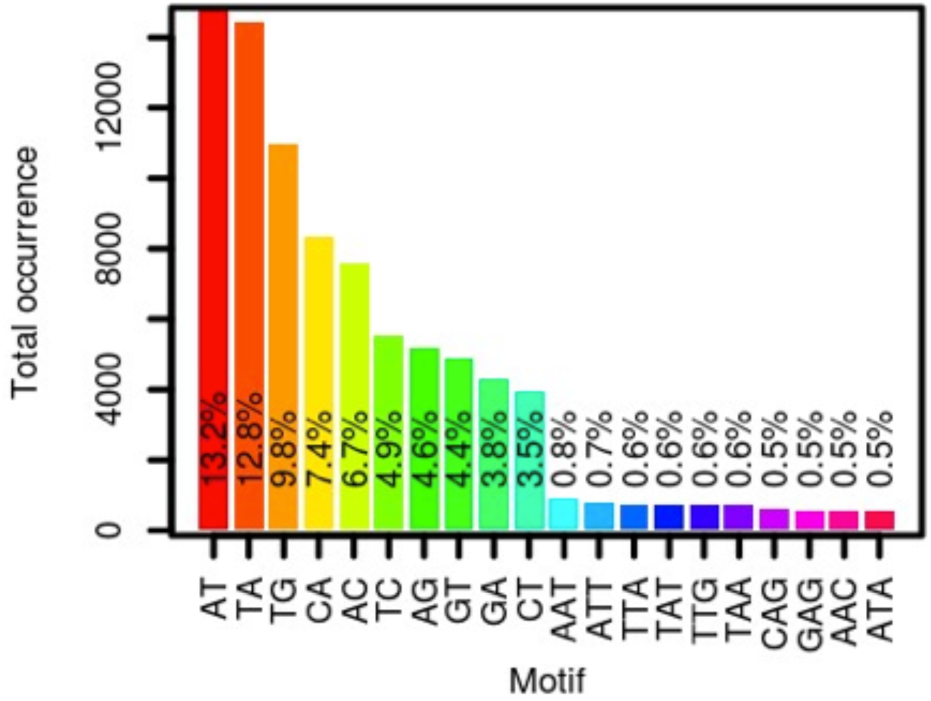
Distribution of the top base-pair composition of microsatellite motifs in the *M. melodia* genome

### Gene annotation and function prediction

The MAKER genome annotation pipeline predicted 15,086 genes and 139 pseudogenes in the song sparrow genome, fewer than *T. guttata, F. albicollis*, and *M. vitellinus*, but higher than *G. fortis* (Table 3). The average gene length, exon length, intron length, and the total number of exons and introns predicted are also less compared to closely related species (Table 3). Out of the 15,086 predicted genes, 12,541 genes were assigned putative functions with BLASTP (File S19). InterProScan assigned functional domains to 11,298 (74.9%) predicted genes (File S20). A total of 7,010 genes obtained GO annotations. Pathway annotations were assigned to 2,716 genes.

**Table 3:** Numbers of genes predicted in the *M. melodia* genome compared to *Taeniopygia guttata* (zebra finch) and *Ficedulla albicollis* (collared flycatcher), *Manacus vitellinus* (golden-collared manakin) and *Geospiza fortis* (medium ground finch).

|  | *M. melodia* | *T. guttata*<sup>1</sup> | *F. albicollis*<sup>2</sup> | *M. vitellinus*<sup>3</sup> | *G. fortis*<sup>4</sup> |
| --- | --- | --- | --- | --- | --- |
| Number of genes | 15,086 | 17,561 | 16,763 | 18,976 | 14,388 |
| Mean gene length (bp) | 14,457 | 26,458 | 31,394 | 27,847 | 30,164 |
| Mean CDS length (bp) | 1,325 | 1,677 | 1,942 | 1,929 | 1,766 |
| Number of exons | 131,940 | 171,767 | 189,043 | 190,390 | 164,721 |
| Mean exon length (bp) | 153 | 225 | 253 | 264 | 195 |
| Mean number of<br>exons/gene | 8.67 | 10.25 | 12.22 | 11.51 | 11.41 |
| Number of introns | 116,724 | 153,909 | 171,236 | 171,089 | 149,563 |
| Mean intron length<br>(bp) | 1,695 | 2,930 | 3,257 | 3,294 | 2,813 |

<sup>1</sup>https://www.ncbi.nlm.nih.gov/genome/annotation_euk/Taeniopygia_guttata/103/
<sup>2</sup>https://www.ncbi.nlm.nih.gov/genome/annotation_euk/Ficedula_albicollis/101/
<sup>3</sup>https://www.ncbi.nlm.nih.gov/genome/annotation_euk/Manacus_vitellinus/102/
<sup>4</sup>https://www.ncbi.nlm.nih.gov/genome/annotation_euk/Geospiza_fortis/101/

Annotated genes were assigned annotation edit distance (AED) scores with values ranging from 0 to 1. AED is a distance metric score that signifies how closely gene models match transcript and protein evidence. Gene models with AED scores closer to 0 have better alignment with the evidence provided in the MAKER pipeline. A distribution of the percentage of genes with their corresponding AED scores shows close similarity of the annotated genes with the transcript and protein evidence provided in the MAKER pipeline (Fig S1).

The song sparrow genome assembly contained 4,318 complete universal single-copy orthologs (BUSCOs; 87.9%) from a total of 4,915 BUSCO groups searched. Among all complete BUSCOs, 99.4% were present as single-copy genes and 0.6% were duplicated. About 7.4% (356) of the orthologous gene models were partially recovered, and 4.9% (241) had no significant matches. The incomplete and missing gene models could either be partially present or missing, or could indicate genes that are too divergent or have very complex structures, making their prediction difficult. Incomplete and missing gene models could also suggest problems associated with the genome assembly and gene annotation. The results from the BUSCO analysis are in agreement with the Circos plot (Fig 5), in which few scaffolds in the zebra finch genome assembly are not represented in our assembly and very few small translocations exist between the two genome assemblies.

**Figure 5:**
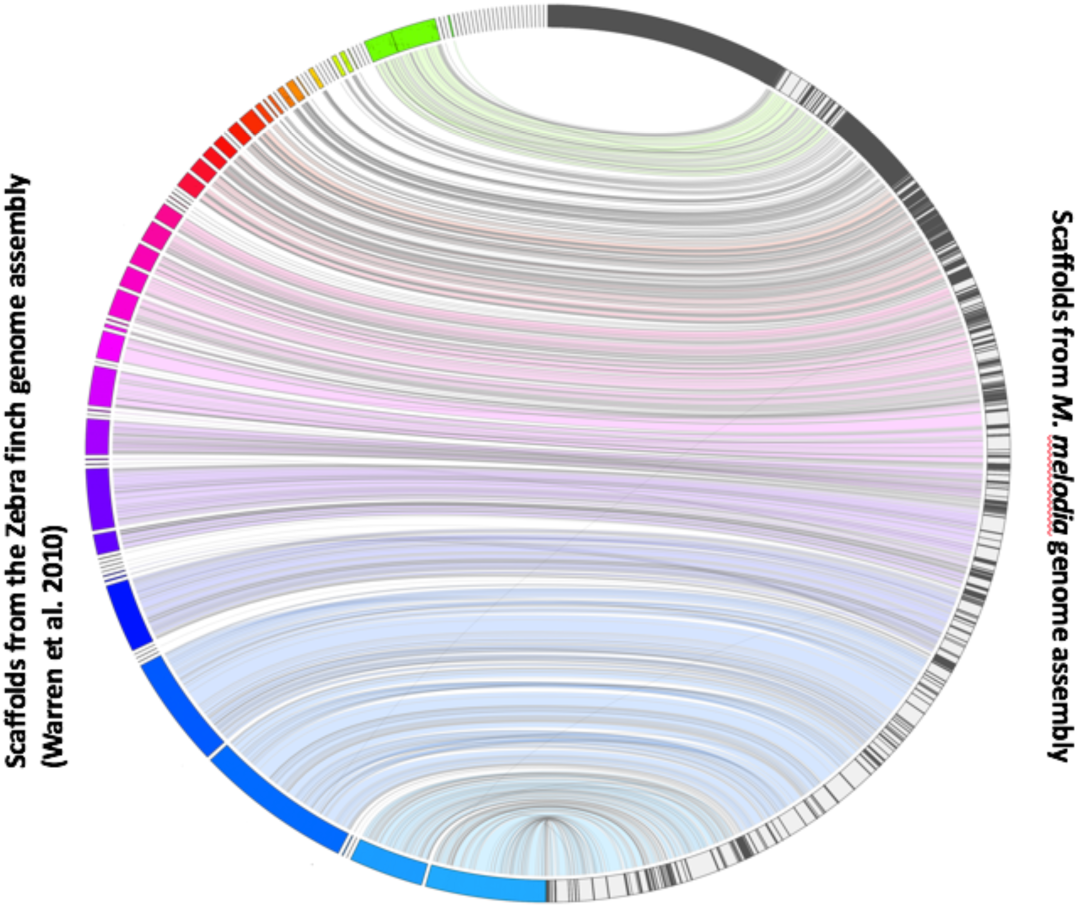
Jupiter plot correlating zebra finch and song sparrow genome assemblies, considering scaffolds greater than 100 kbp in the reference zebra finch genome and the largest scaffolds representing 95% of the query song sparrow genome.

### Non-coding RNA prediction and identification

A total of 267 tRNAs were detected in the song sparrow genome by tRNAscan-SE (see File S21 for sequence and structure of tRNAs), out of which 129 were found coding for the standard twenty amino acids. The predicted output from tRNAscan-SE (File S22) contained 114 tRNAs with low Infernal as well as Isotype scores; these were characterized as pseudogenes lacking tRNA-like secondary structures (Lowe and Chan 2016). Two tRNAs had undetermined isotypes and 22 were chimeric, with mismatch isotypes. Chimeric tRNAs contain point mutations in their anticodon sequence, rendering different predicted isotypes than those predicted by structure-specific tRNAscan-SE covariance models. Among all predicted tRNAs, 11 contained introns within their sequences. No suppressor tRNAs and tRNAs coding for selenocysteine were predicted. The subset of 10 randomly selected tRNAs was also predicted in many other species in both GtRNAdb and tRNAdb databases.

Infernal searches predicted a total of 364 ncRNAs in the song sparrow genome, comprising 166 miRNAs, 8 rRNAs, 154 snoRNAs, 16 snRNAs, and 20 lncRNAs (File S23). Compared to the genomes of related avian species (*T. guttata* and *F. albicollis*), the song sparrow genome has the highest number of predicted tRNAs, but fewer other ncRNAs (Table 4).

**Table 4:** Numbers of ncRNAs predicted in the *Melospiza melodia* genome compared to *Taeniopygia guttata* (zebra finch) and *Ficedulla albicollis* (collared flycatcher).

|  | *M. melodia* | *T. guttata*<sup>1,2</sup> | *F. albicollis*<sup>1,3</sup> |
| --- | --- | --- | --- |
| tRNA | 267 | 184 | 179 |
| miRNA | 166 | 302 | 510 |
| snRNA | 16 | 44 | 32 |
| snoRNA | 154 | 241 | 199 |
| rRNA | 8 | 100 | 22 |
| lncRNA | 20 | 908 | 1473 |

<sup>1</sup>http://useast.ensembl.org/info/data/ftp/index.html
<sup>2</sup>https://www.ncbi.nlm.nih.gov/genome/annotation_euk/Taeniopygia_guttata/103/
<sup>3</sup>https://www.ncbi.nlm.nih.gov/genome/annotation_euk/Ficedula_albicollis/101/

## CONCLUSION

The Chicago and shotgun sequencing libraries along with the HiRise assembly software enabled accurate and highly contiguous *de novo* assembly of the song sparrow genome. The genome assembly is 978.3 Mb, with 48 scaffolds (L50) making up half the genome size. A previous estimate of genome size of *M. melodia* from densitometry analysis provided a C-value of 1.43 pg (1,398.54 Mb) (Gregory et al. 2009). Our own *k-mer* based estimate of genome size from paired reads in the shotgun and Chicago libraries using Kmergenie v1.7044 (Chikhi and Medvedev 2013) yielded an estimated size of 1,127.25 Mb. Both these genome size estimates and the genome sizes of related birds (Table S1) are slightly higher than our genome assembly (978.3 Mb); this could be due to the low TE content observed in the song sparrow. Vertebrate genome sizes are typically correlated with TE content and birds are no exception (Kidwell 2002, Gregory and Elliott 2015). Indeed, birds, along with bats, are proposed to have been subject to extensive DNA loss via an ‘accordion model’ of genome evolution (Kapusta et al. 2017). Our data suggest that, among birds, the song sparrow may have been particularly susceptible to the segmental deletions that are critical to this model. The small genome size of the song sparrow can also be attributed to the compression of repetitive regions, which is generally observed in assemblies generated from short-read sequencing data. This is also consistent with the fact that our genome contains fewer repeats when compared to related songbirds (Fig 2). Although short reads limit our ability to characterize all genomic repeats, we have been able to resolve the vast majority of repeats into LINEs, SINEs, LTRs, and DNA retrotransposons (Fig 2, Table 2).

Our genome is highly complete, with 87.5% full-length genes present out of 4,915 universal orthologous genes in avian species. A large set of genes (15,086) with known homology to related birds was annotated in our study. A majority of these genes (83%) were assigned with putative functions. The improved scaffold lengths and gene model annotations will facilitate studies to identify genes responsible for multiple phenotypic traits of interest. Additionally, longer scaffolds in the Chicago HiRise assembly will help detect regions under selection, including SNPs and structural variants such as insertions/deletions or copy number variations which are potentially responsible for the phenotypic diversity observed in this species.

Though we report fewer miRNAs, snRNAs, snoRNAs, rRNAs, and lncRNAs in this genome than in related songbirds, we have high confidence in the predicted ncRNAs because we used conservative cutoffs to reduce false positives. Pending the availability of long-read data, this genome assembly provides an excellent reference for a range of genetic, ecological, functional, and comparative genomic studies in song sparrows and other songbirds.

## ACKNOWLEDGEMENTS

This research was supported in part by an Institutional Development Award (IDeA) from the National Institute of General Medical Sciences of the NIH (P20GM103395). The content is solely the responsibility of the authors and does not necessarily reflect the official views of the NIH. We also thank David Sonneborn and Jack Withrow for their roles in making a vouchered specimen available for our work. We thank the staff of Dovetail Genomics for help in preparing and processing the Chicago library and HiRise assemblies. The high performance computing cluster at Georgia Advanced Computing Resource Center (GACRC) at the University of Georgia provided computational infrastructure and technical support throughout the work.

