## Supplementary File S2 for "A high-quality genome assembly of the North American Song Sparrow, *Melospiza melodia*"

[set]:gmat
#1. gmat.pl-------------------------------------------------------------------------------------------
# Parameter "run" takes the value "Y" or "N". This module will be run if "run" equals to value "Y". Otherwise , This module will be ignored.   
# e.g. ModulRun = Y or ModulRun = N. Default is Y.
ModulRun = Y

#other parameters:
#-i =
# NECESSARY, Input SequenceFile.
-r = 5
# Minimum repeated times (Repeat-times) of motif, integer value, >0, the default is 5.
-m = 2
# Minimum length (Mini-length) of motif, integer value, >0, the default is 2.
-x = 20
# Maximum length (Maxi-length) of motif, the default is 10, value of x greater or equal to value of 
# m, if value x= value m, mining motif at only this given length, meaning one length of motif.
-s = 0
# Highlighting repeat sequence or not, the value should be 0 or 1.  
# 1 meaning displaying original DNA sequence in lowcase and repeated sequence in upcase;
# 0 meaning no flanking sequence output in SSR loci file.

[set]:ssrfig

#2. ssrfig.pl-----------------------------------------------------------------------------------------
# Parameter "run" takes the value "Y" or "N". This module will be run if "run" equals to value "Y". Otherwise , This module will be ignored.   
# e.g. ModulRun = Y or ModulRun = N. Default is Y.
ModulRun = Y

#other parameters:
# -i =  
# Input File (e.g. -i = SequenceFileName.ssr.sat2). Default is SequenceFileName.ssr.sat2 created by 
# gmat.pl, if HERE is NOT SPECIFIED (left empty).
-f1 = 20
# The number of top k-mers to be displayed in bar chart (integer value). Default value is 10.
-f2 = 20
# The number of top motifs to be displayed in bar chart (integer value). Default value is 20.
-f3 = 20
# The number of top grouped motifs to be displayed in bar chart (integer value). Default value is 20.
-f4 = 20
# The number of top SSR loci on chromosomes to be displayed in bar chart (integer value). Default 
# value is 10.
-f5 = 20
# The number of top distribution of SSR length to be displayed in bar chart (integer value). Default 
# value is 20.

[set]:ssr2gff
#3.1 ssr2gff.pl----------------------------------------------------------------------------------------
# Parameter "run" takes the value "Y" or "N". This module will be run if "run" equals to value "Y". Otherwise , This module will be ignored.   
# e.g. ModulRun = Y or ModulRun = N. Default is Y.
ModulRun = N

#other parameters:
# -i = 
# Input SequenceFile (e.g. -i = SequenceFileName.ssr). Default is SequenceFileName.ssr produced by gmat.pl, if 
# HERE is NOT SPECIFIED (left empty).

[set]:ssr2gff3
#3.2 ssr2gff3.pl---------------------------------------------------------------------------------------
# following options won't work unless THIS VALUE is Y
ModulRun = Y

#-i = 
# Input SequenceFile (e.g. -i = SequenceFileName.ssr). Default is SequenceFileName.ssr produced by gmat.pl, if 
# HERE is NOT SPECIFIED (left empty).

[set]:doprimer_smt
#4. doprimer_smt.pl-----------------------------------------------------------------------------------
# Parameter "run" takes the value "Y" or "N". This module will be run if "run" equals to value "Y". Otherwise , This module will be ignored.   
# e.g. ModulRun = Y or ModulRun = N. Default is Y.
ModulRun = N

#other parameters:
#-i = 
# Input SequenceFile (e.g. -i = SequenceFileName). Default is SequenceFileName used in gmat.pl, if 
# HERE is NOT SPECIFIED (left empty).
#-sr = 
# Input SSRFile (e.g. –sr = SequenceFileName.ssr). Default is SequenceFileName.ssr created by gmat.pl
# if HERE is NOT SPECIFIED (left empty).
-p = w
# Platform, the value can only be u or w (u: Unix, w: windows).Default is w.
-fl = 200
# Extracted SSR flanking sequence length at either side of SSR loci(integer value). default value is 200.
-l = 2000
# Length of maximum allowed template for primer designing (integer value), default value is 2000.
-smin = 120
# Minimum length of amplicon size (integer value), default value is 100.
-Smax = 400
# Maximum length of amplicon size (integer value), default value is 400.
-tm = 60
# Optimal annealing temperature for PCR primer (integer value), default value is 60
#-dir=
# Directory or path of primer3_core program. The actual directory should be given. if path is set up in envroment variable, keep this unchanged. note: no "/" in the end.e.g. -dir=/home/SCE/wangxuewen/bin/primer3-2.2.3/src.
-ts = F
# Logic T or F, Output Template sequence with primer (T) or not(F). default is F.
-n = MK
#the prefix name of the markers. User can give any prefix name. Default is "MK". e.g. MK1, MK200

[set]:elctPCR
#5. elctPCR.pl----------------------------------------------------------------------------------------
# Parameter "run" takes the value "Y" or "N". This module will be run if "run" equals to value "Y". Otherwise , This module will be ignored.   
# e.g. ModulRun = Y or ModulRun = N. Default is Y.
# Ignore the message prompted as "WARNING: 1sts have incomplete description line" in DOS platform. This will not affect the results. 
ModulRun = N

#other parameters
-i = 
# Input SequenceFile (e.g. -i = SequenceFileName) for markers to be mapped to.  If another # sequences file is given here, the markers 
# will be mapped to the new # sequences file (e.g. -i = anotherseq.fa). 
# Otherwise, leave the value empty as is and then the markers will be mapped to the sequences where the markers originated if HERE is NOT SPECIFIED (empty).

#-mk = 
# Input MarkerFile (e.g. –mk = SequenceFileName.ssr.finalPrimer.txt.sts). Default is 
# SequenceFileName.ssr.finalPrimer.txt.sts created by doprimer_smt.pl, if HERE is NOT SPECIFIED 
# (left empty).
-o = eMap
# eMapping tag of output files. default tag is "eMap". e.g. mapping result file is sequence file name
# plus ".eMap". There are four files will produced. There are $seq.eMap, $seq.eMap.amp, 
# $seq.eMap.frg, $seq.eMap.frg.sat4.
-pf = w
# System platform, the value can only be u or w (u: Unix, w: windows).
-w = 12
# Word size for ePCR, default is 12. For more details, refer to e-PCR software.
-f = 1
# value takes 1 or 3. 1 for contiguous words and 3 for discontiguous.  Default value is 1. For more 
# details, refer to e-PCR software.
-m = 3000
# Margin. default is 3000. For more details, refer to e-PCR software.
-d = 100-1000
# Set allowed sts size range: value1-value2, e.g. 100-1000. default is 100-1000. For more details, refer to e-PCR software.
-n = 0
# Max mismatches allowed (integer value, default 0, suggested value is 1). For more details, refer 
#to e-PCR software.
-g = 0
# Max indels allowed (integer value, default 0, suggested value is 1). For more details, refer to 
#e-PCR software.
-p = -
# value is -|+. Turn hits post-process on/ for e-PCR. Default is -. For more details, refer to e-PCR 
# software.

[set]:mk2gff3
#5. mk2gff3.pl----------------------------------------------------------------------------------------
# Parameter "run" takes the value "Y" or "N". This module will be run if "run" equals to value "Y". Otherwise , This module will be ignored.   
# e.g. ModulRun = Y or ModulRun = N. Default is Y.
# Ignore the message prompted as "WARNING: 1sts have incomplete description line" in DOS platform. This will not affect the results. 
ModulRun = N

#other parameters:
-o = eMap
# This value usually should be the same value of -o tag used in [sect] elctPCR. The value is the tag of file with suffix ".frg", that is the content of xxx letters of in last part of file name "[prefix].xxx.frg".  If mapped to the same genome which markers are designed from, just leave the -o tag value as default "eMap".  With this information, the Gbrowse will display all alleles information of the markers when mapped to -o tag genome sequences. It is useful when markers are designed from specie one and want to show the amplicon size in specie two in Gbrowse. Actually the mapping result called .frg already has the data. Here is just to show in Gbowse. 
