## Supplementary File S6 for "A high-quality genome assembly of the North American Song Sparrow, *Melospiza melodia*"

#-----Genome (these are always required)
genome=/scratch/sl50708/songsparrow/genemark/song_sparrow_04Oct2016_v2yLs.fasta #genome sequence (fasta file or fasta embeded in GFF3 file)
organism_type=eukaryotic #eukaryotic or prokaryotic. Default is eukaryotic

#-----Re-annotation Using MAKER Derived GFF3
maker_gff= #MAKER derived GFF3 file
est_pass=0 #use ESTs in maker_gff: 1 = yes, 0 = no
altest_pass=0 #use alternate organism ESTs in maker_gff: 1 = yes, 0 = no
protein_pass=0 #use protein alignments in maker_gff: 1 = yes, 0 = no
rm_pass=0 #use repeats in maker_gff: 1 = yes, 0 = no
model_pass=0 #use gene models in maker_gff: 1 = yes, 0 = no
pred_pass=0 #use ab-initio predictions in maker_gff: 1 = yes, 0 = no
other_pass=0 #passthrough anyything else in maker_gff: 1 = yes, 0 = no

#-----EST Evidence (for best results provide a file for at least one)
est=/scratch/sl50708/songsparrow/evidence/transcriptome/melospiza_transcriptome.fasta  #set of ESTs or assembled mRNA-seq in fasta format
altest=/scratch/sl50708/songsparrow/evidence/transcriptome/Junco_transcriptome.fasta #EST/cDNA sequence file in fasta format from an alternate organism
est_gff= #aligned ESTs or mRNA-seq from an external GFF3 file
altest_gff= #aligned ESTs from a closly relate species in GFF3 format

#-----Protein Homology Evidence (for best results provide a file for at least one)
protein=/scratch/sl50708/songsparrow/evidence/protein_data/Melospiza_Taeniopygia_Junco.fasta  #protein sequence file in fasta format (i.e. from mutiple oransisms)
protein_gff=  #aligned protein homology evidence from an external GFF3 file

#-----Repeat Masking (leave values blank to skip repeat masking)
model_org=  #select a model organism for RepBase masking in RepeatMasker
rmlib=/scratch/sl50708/song2/maker/aves_plus_10-17-2018.fasta  #provide an organism specific repeat library in fasta format for RepeatMasker
repeat_protein=/db/maker/2.31.9/data/te_proteins.fasta #provide a fasta file of transposable element proteins for RepeatRunner
rm_gff= #pre-identified repeat elements from an external GFF3 file
prok_rm=0 #forces MAKER to repeatmask prokaryotes (no reason to change this), 1 = yes, 0 = no
softmask=1 #use soft-masking rather than hard-masking in BLAST (i.e. seg and dust filtering)

#-----Gene Prediction
snaphmm=/scratch/sl50708/song2/snap_round2/param/snap_songsparrow.hmm  #SNAP HMM file
gmhmm= #GeneMark HMM file
augustus_species=songsparrow  #Augustus gene prediction species model
fgenesh_par_file= #FGENESH parameter file
pred_gff= #ab-initio predictions from an external GFF3 file
model_gff= #annotated gene models from an external GFF3 file (annotation pass-through)
est2genome=0  #infer gene predictions directly from ESTs, 1 = yes, 0 = no
protein2genome=0  #infer predictions from protein homology, 1 = yes, 0 = no
trna=0 #find tRNAs with tRNAscan, 1 = yes, 0 = no
snoscan_rrna= #rRNA file to have Snoscan find snoRNAs
unmask=0 #also run ab-initio prediction programs on unmasked sequence, 1 = yes, 0 = no

#-----Other Annotation Feature Types (features MAKER doesn't recognize)
other_gff= #extra features to pass-through to final MAKER generated GFF3 file

#-----External Application Behavior Options
alt_peptide=C #amino acid used to replace non-standard amino acids in BLAST databases
cpus=1 #max number of cpus to use in BLAST and RepeatMasker (not for MPI, leave 1 when using MPI)

#-----MAKER Behavior Options
max_dna_len=100000 #length for dividing up contigs into chunks (increases/decreases memory usage)
min_contig=1 #skip genome contigs below this length (under 10kb are often useless)

pred_flank=200 #flank for extending evidence clusters sent to gene predictors
pred_stats=0 #report AED and QI statistics for all predictions as well as models
AED_threshold=1 #Maximum Annotation Edit Distance allowed (bound by 0 and 1)
min_protein=0 #require at least this many amino acids in predicted proteins
alt_splice=0 #Take extra steps to try and find alternative splicing, 1 = yes, 0 = no
always_complete=0 #extra steps to force start and stop codons, 1 = yes, 0 = no
map_forward=0 #map names and attributes forward from old GFF3 genes, 1 = yes, 0 = no
keep_preds=0 #Concordance threshold to add unsupported gene prediction (bound by 0 and 1)

split_hit=10000 #length for the splitting of hits (expected max intron size for evidence alignments)
single_exon=0 #consider single exon EST evidence when generating annotations, 1 = yes, 0 = no
single_length=250 #min length required for single exon ESTs if 'single_exon is enabled'
correct_est_fusion=0 #limits use of ESTs in annotation to avoid fusion genes

tries=2 #number of times to try a contig if there is a failure for some reason
clean_try=0 #remove all data from previous run before retrying, 1 = yes, 0 = no
clean_up=0 #removes theVoid directory with individual analysis files, 1 = yes, 0 = no
TMP= #specify a directory other than the system default temporary directory for temporary files
