## Supplementary figures and images for "A high-quality genome assembly of the North American Song Sparrow, *Melospiza melodia*"

### Supplementary Fig S1

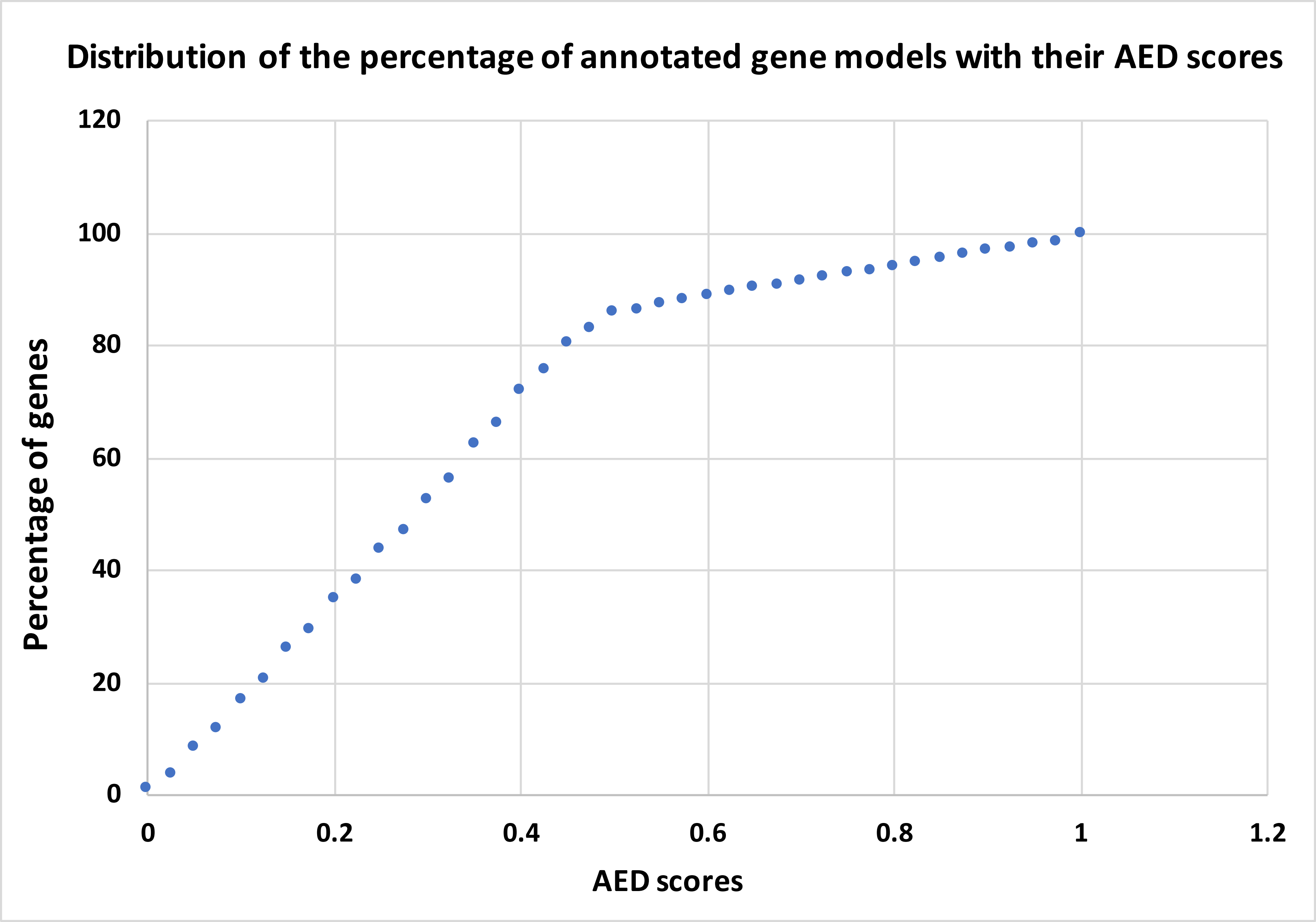
